# Whole-Genome Sequencing Uncovers Chromosomal and Plasmid-Borne Multidrug Resistance and Virulence Genes in Poultry-Associated *Escherichia coli* from Nigeria

**DOI:** 10.1101/2025.09.18.677015

**Authors:** Chika P. Ejikeugwu, Chijioke Edeh, Emmanuel A. Nwakaeze, Michael U. Adikwu, Carmen Torres, Christopher J. Creevey, Peter M. Eze

## Abstract

**Background:** Unregulated antibiotic use in poultry farming drives the emergence of multidrug-resistant (MDR) bacteria, which can spread to humans through the food chain and environment, posing serious public health risks. Whole-genome sequencing (WGS), combined with antimicrobial susceptibility testing (AST), enables detailed characterization of resistance mechanisms and supports antimicrobial stewardship. This study investigated the phenotypic and genotypic antimicrobial resistance (AMR), plasmid content, and virulence factors of an MDR *Escherichia coli* strain isolated from chicken droppings in Enugu State, Nigeria.

**Results:** Disk diffusion AST showed resistance to six of seven antibiotics tested (cefotaxime, ampicillin, erythromycin, gentamicin, ciprofloxacin, and doxycycline). Broth microdilution confirmed elevated minimum inhibitory concentrations across multiple classes, indicating an MDR phenotype. Hybrid WGS (Illumina and Nanopore) produced a 5.33 Mb genome comprising one chromosome and four plasmid-associated contigs. Chromosomal antimicrobial resistance genes (ARGs), including *aac(6’)-Ib-cr, bla*_CTX-M-15_, and *bla*_OXA-1_, conferred resistance to aminoglycosides, fluoroquinolones, cephalosporins, and penicillins. Additional resistance determinants included efflux pumps, transport-associated genes, regulatory elements, and membrane modification genes. Plasmid-borne ARGs conferring resistance to aminoglycosides, trimethoprim, macrolides, sulfonamides, penicillins, and tetracyclines were also identified. The presence of Col156 and IncF-type plasmids indicates strong potential for horizontal gene transfer. Virulence profiling revealed numerous chromosomally encoded factors related to adhesion, iron acquisition, and toxin production. These included the *pap* gene cluster encoding P fimbriae; adhesion-associated genes (*yagW/ecpD, ykgK/ecpR*); multiple iron acquisition systems (enterobactin, yersiniabactin, aerobactin, and heme uptake); and *sat*, which encodes an autotransporter toxin. Additionally, the plasmid-borne gene *senB* encodes an enterotoxin that induces intestinal fluid secretion and contributes to diarrheal disease. Multilocus sequence typing identified the strain as ST131, a globally disseminated high-risk lineage.

**Conclusions:** This study provides a comprehensive genomic characterization of an MDR *E. coli* strain from poultry, revealing multiple chromosomal and plasmid-borne ARGs, and a diverse virulence gene repertoire. The detection of ST131 in poultry waste highlights a complex public health issue involving zoonotic transmission, veterinary impact, and environmental spread of AMR. These findings underscore the need for prudent antibiotic use, continuous monitoring, and integrated genomic surveillance across agricultural and environmental sectors, alongside improved antimicrobial stewardship and strengthened farm biosecurity measures.

## INTRODUCTION

The growing threat of antimicrobial resistance (AMR) is closely tied to the extensive use of antibiotics in food-animal production. In poultry farming, antibiotics are frequently employed for prophylaxis, growth promotion, and treatment, often without adequate regulation or oversight [1,2]. This practice imposes a strong selective pressure that fosters the emergence and persistence of multidrug-resistant (MDR) bacteria, which can be transmitted to humans through food, direct contact, and environmental routes [3]. Within a One Health framework recognizing the interconnectedness of human, animal, and environmental health, such transmission pathways underscore the broader public health significance of antimicrobial use in livestock systems.

Among Gram-negative bacteria, *Escherichia coli* plays a pivotal role in the AMR crisis. While many strains are harmless commensals, pathogenic *E. coli* is broadly classified into intestinal (InPEC) and extraintestinal (ExPEC) pathotypes. Intestinal pathotypes include enterohemorrhagic (EHEC), enterotoxigenic (ETEC), enteropathogenic (EPEC), enteroaggregative (EAEC), and enteroinvasive (EIEC) *E. coli*, which are associated with a spectrum of diarrheal diseases. In contrast, extraintestinal pathogenic *E. coli* (ExPEC) are responsible for infections outside the gut, including urinary tract infections, neonatal meningitis, and sepsis [4,5]. *E. coli* exhibits remarkable genetic adaptability and an exceptional capacity for acquiring antibiotic resistance genes (ARGs), particularly through horizontal gene transfer (HGT) mechanisms [6,7]. The global proliferation of extended-spectrum β-lactamase (ESBL) and metallo-β-lactamase (MBL) producing *E. coli* strains has rendered many frontline antibiotics such as cephalosporins and carbapenems ineffective [8,9], prompting the World Health Organization (WHO) to classify *E. coli* as a critical priority pathogen [10]. Poultry farms serve as significant reservoirs for resistant *E. coli*, given the organism’s ubiquity in avian gastrointestinal tracts and its genomic plasticity [6,11].

In Nigeria, where the poultry industry has rapidly expanded, antibiotic use remains largely unregulated, contributing to the selection and amplification of MDR bacteria [9,12]. Previous studies in southwestern Nigeria have revealed high resistance levels in *E. coli* isolates from poultry environments, including resistance to tetracycline (81%), sulfamethoxazole (67%), and streptomycin (56%), with 85% of isolates carrying the ^*bla*^TEM gene and 14% harboring class 1 integrons [13]. The role of poultry farms as hotspots for ARG dissemination is further underscored by the detection of resistant *E. coli* in abattoirs, water bodies, and soil surrounding poultry operations [2,14,15,16,17).

Furthermore, ESBL-producing *E. coli* strains frequently co-occur with other MDR pathogens such as *Klebsiella pneumoniae* and *Pseudomonas aeruginosa*, increasing the risk of cross-species gene exchange [9,18,19,20]. Given the high mobility of plasmid-encoded ARGs, poultry operations may act as key nodes in the wider transmission network of resistance determinants [21,22]. Human exposure to resistant *E. coli* from poultry can occur through multiple pathways. Contaminated meat, particularly when improperly handled or undercooked, represents a major source of transmission [23]. Farm workers and residents are also at heightened risk through occupational contact [24,25]. Additionally, environmental runoff from poultry waste introduces ARGs and resistant bacteria into surrounding ecosystems, contaminating water, soil, crops irrigated with polluted water, and the general environment [9,26,27].

Once these resistant strains establish in human populations, they contribute to a growing burden of difficult-to-treat infections, leading to increased hospitalization, healthcare costs, and mortality. The threat is exacerbated by the global spread of mobile colistin resistance (*mcr*) genes, which enable transferable resistance to colistin, a last-resort antibiotic [28]. Recognized as a critical concern by the World Health Organization, their rapid horizontal dissemination threatens treatment efficacy, alongside the declining effectiveness of other last-resort antibiotics such as carbapenems [8,29]. It has been presumed that by 2050, AMR could claim 10 million lives annually if left unchecked [8]. To mitigate the public health impact of poultry-associated AMR, robust surveillance and molecular characterization of resistant strains are essential, particularly in resource-limited settings. High-throughput genomic tools, especially whole-genome sequencing (WGS), provide a powerful means to identify resistance determinants, infer transmission pathways, and understand the evolutionary dynamics of MDR *E. coli* strains [30,31].

In regions such as Nigeria, where regulatory frameworks remain underdeveloped, the application of WGS could offer critical insights to guide evidence-based policy and intervention strategies [32]. Efforts to curb AMR spread in poultry production must also include improved biosecurity, reduced antibiotic use, and the promotion of vaccination-based disease control [32]. Strengthening antimicrobial stewardship and enforcing guidelines on antibiotic use in livestock are urgent priorities for Nigeria and other sub-Saharan African nations [32,33].

In our previous study surveying poultry farms in Enugu State, Nigeria, we found that 90.5% of farmers routinely used antibiotics, yet 65% were unaware of AMR, and only 16% recognized its potential health risks [32]. This highlights critical gaps in antibiotic stewardship and underscores the potential for the selection and dissemination of MDR pathogens in poultry systems and the broader environment. From this earlier investigation, we isolated an MDR *E. coli* strain with an interesting antibiotic-resistance profile from one of the surveyed farms. In this paper, we report the phenotypic and genotypic characterization of the isolate’s chromosomal and plasmid-mediated resistome, as well as its virulence genes, with the aim of linking the observed resistance patterns to the broader issue of antibiotic misuse in poultry farming.

## MATERIALS AND METHODS

### Isolation of Bacterium and MALDI-TOF Identification

A bacterium was isolated from chicken droppings from a poultry farm in Enugu State, Nigeria following standard microbiological procedure for bacterial isolation [33]. The poultry farm was one of the farms previously surveyed for antibiotic use and antimicrobial resistance (AMR) awareness in Enugu State, Nigeria reported in our previous study [32]. The isolate was subjected to preliminary biochemical identification, including culture in Coliform *ChromoSelect* Agar (CCA; Sigma-Aldrich, Germany). A fully automated microbial mass spectrometry detection system (Autof MS1600 MALDI-TOF, Autobio Labtec Instrument Co., Ltd., China) was also used to confirm the isolate’s identity following a previously described method [34].

### Antibiotic Susceptibility Testing (AST)

AST of the isolated bacterium was conducted using the standard disk diffusion method, according to our previously described protocol [9], and in compliance with the guidelines of the European Committee on Antimicrobial Susceptibility Testing (EUCAST) [35] and the Clinical and Laboratory Standards Institute (CLSI) [36]. A volume of 20 mL of molten Mueller-Hinton (MH) agar (Sigma-Aldrich, India) was poured into 90 mm Petri plates and allowed to solidify. The isolate, previously grown overnight in MH broth, was adjusted to a 0.5 McFarland standard (1 × 10^8^ CFU/mL) and uniformly spread on the agar plates using a sterile swab stick. Antibiotic discs (Oxoid, UK) containing cefotaxime (30 µg), ampicillin (10 µg), erythromycin (15 µg), gentamicin (10 µg), ciprofloxacin (5 µg), doxycycline (30 µg), and imipenem (10 µg) were applied at 20 mm intervals. Plates were incubated at 37°C for 18-24 h, after which inhibition zone diameters (IZDs) were measured in mm. The experiment was performed in triplicate, and the results were interpreted according to the EUCAST inhibition zone diameter breakpoints using the antibiotic-susceptible reference strain *E. coli* ATCC 25922 [35].

### Determination of Minimum Inhibitory Concentrations (MIC)

The minimum inhibitory concentrations (MICs) of 10 antibiotics representing major clinical classes were determined using the standard broth microdilution method in cation-adjusted MH broth (CAMHB; Sigma-Aldrich, India). The procedure followed established protocols [34] and complied with EUCAST and CLSI guidelines [37,38]. The antibiotics tested included gentamicin sulfate salt (aminoglycoside; Sigma-Aldrich, China), ciprofloxacin (fluoroquinolone; Merck, USA), imipenem monohydrate and meropenem trihydrate (carbapenems; Glentham Life Sciences Ltd, UK), cephalexin (cephalosporin; Cayman Chemical Company, USA), tetracycline (tetracycline; Sigma-Aldrich, Israel), amoxicillin and amoxicillin trihydrate:potassium clavulanate (4:1) (penicillins; Sigma-Aldrich, China and Israel, respectively), colistin sulfate salt (polymyxin; Sigma-Aldrich, China), and erythromycin (macrolide; Sigma-Aldrich, China). Antibiotics were dissolved in sterile CAMHB to achieve a stock concentration of 200 µg/mL and were serially diluted (two-fold) in sterile, round-bottom 96-well polystyrene microplates to 0.006 µg/mL. An equal volume of standardized bacterial inoculum was added to achieve final antibiotic concentrations ranging from 100 µg/mL to 0.003 µg/mL and a cell density of 1 × 10^6^ CFU/mL. Plates were incubated at 37°C for 18 h, and growth inhibition was assessed visually based on medium turbidity. MICs were defined as the lowest concentrations that completely inhibited visible bacterial growth. All assays were performed in triplicate, and results were interpreted using EUCAST breakpoints, with the antibiotic-susceptible reference strain *E. coli* ATCC 25922 as a control [35].

### Whole-genome Sequencing, Assembly, Annotation and Isolate’s identity Confirmation by 16S Taxonomy

Genomic DNA extraction and hybrid sequencing using Illumina and Nanopore platforms were performed by MicrobesNG (Birmingham, UK) following standard protocols (https://microbesng.com/), and as previously described [34]. Hybrid genome assembly was conducted using Unicycler v0.4.0, and assembly quality was assessed with QUAST via the Galaxy platform (https://usegalaxy.org/). Genome annotation was performed using Prokka v1.14.6. GC content for the complete genome and individual contigs was calculated using the GC Content Calculator (https://jamiemcgowan.ie/bioinf/gccontent.html). The 16S rRNA gene was extracted using Extractseq on Galaxy and used for species confirmation via BLAST searches against the NCBI rRNA/ITS databases (https://blast.ncbi.nlm.nih.gov/Blast.cgi). The sequence was subsequently submitted to GenBank. Molecular typing was performed by multilocus sequence typing (MLST) using MLST v2.32.2 through starAMR v0.12.1 on Galaxy, with reference to PubMLST schemes [39].

### *In silico* characterization of antimicrobial resistance genes (ARGs), plasmid replicons, and virulence factors

Assembled contigs were analyzed to identify ARGs using AMRFinderPlus v3.12.8 and starAMR v0.12.1 (which searched against the ResFinder, PlasmidFinder, and PointFinder databases), both available on Galaxy [40]. Additional analysis was performed with the Resistance Gene Identifier (RGI v6.0.5) from the Comprehensive Antibiotic Resistance Database (CARD v4.0.1) (https://card.mcmaster.ca/analyze/rgi) [41]. These tools detect acquired resistance genes, virulence factors, and relevant mutations using curated reference databases and hidden Markov models. Plasmid replicons were predicted using PlasmidFinder v2.1.6 [42], MOB-suite MOB-recon v3.1.9 [43], and starAMR v0.12.1 [40], all accessed via Galaxy. Virulence factors were identified using the Virulence Factor Database through ABRicate v1.0.1, available on Galaxy [44]. Analysis thresholds were set at a minimum DNA identity of 99% and minimum coverage of 80%.

## RESULTS

### Recovery and Characterization of *Escherichia coli* from Poultry Droppings

A bacterial isolate was recovered from chicken droppings collected at a poultry farm in Enugu State, Nigeria. On Coliform *ChromoSelect* Agar (CCA), the isolate formed blue colonies characteristic of *E. coli* (Fig. S1, supplementary material). Identification was further supported by standard cultural and biochemical characterizations and finally confirmed using matrix assisted laser desorption ionization time-of-flight (MALDI-TOF) mass spectrometry. The isolate was designated *E. coli* strain S3 and selected for antimicrobial susceptibility profiling and downstream molecular biology.

### Extensive AMR pattern in *E. coli* Strain S3 Revealed by Disc Diffusion

Compared to the EUCAST breakpoints for the susceptible reference strain *E. coli* ATCC 25922, the IZDs recorded for *E. coli* strain S3 showed resistance to all seven tested antibiotics by disc diffusion AST. The strain exhibited complete resistance (IZD of 0 mm) to six of the seven antibiotics: cefotaxime, ampicillin, erythromycin, gentamicin, ciprofloxacin, and doxycycline. Only imipenem (10 μg) showed a measurable IZD of 7.3 ± 0.6 mm, which is well below the EUCAST breakpoint for *E. coli* ATCC 25922 (26–32 mm). These results indicate broad resistance across β-lactams, aminoglycosides, fluoroquinolones, and macrolides. Consequently, *E. coli* strain S3 was classified as MDR.

### High-Level Resistance Confirmed by Broth Microdilution: MIC Assessment

MIC testing confirmed multidrug resistance in the strain. Gentamicin, ciprofloxacin, tetracycline, and erythromycin each exhibited MICs ≥100 µg/mL (Table 1), far exceeding clinical breakpoints. Similarly, β-lactams including cephalexin, amoxicillin, and amoxicillin-clavulanate showed elevated MICs (50 to ≥100 µg/mL). In contrast, the strain remained susceptible to carbapenems and colistin, with low MICs for imipenem (0.19 µg/mL), meropenem (0.024 µg/mL), and colistin (0.19 µg/mL), indicating preserved efficacy of last-resort antimicrobials.

**Table 1:** Phenotypic AMR profile of *E. coli* strain S3.

| Antibiotics | Class | Inhibition zone diameter [IZD (mm)] |  |
| --- | --- | --- | --- |
|  |  | *E. coli* strain S3 | EUCAST Breakpoint for *E. coli* ATCC 25922 [35] |
| Cefotaxime (30 µg) | Beta-lactam | 0.0 ± 0.0 | 25-31 |
| Ampicillin (10 µg) | Beta-lactam | 0.0 ± 0.0 | 15-22 |
| Erythromycin (15 µg) | Macrolide | 0.0 ± 0.0 | - |
| Gentamicin (10 µg) | Aminoglycoside | 0.0 ± 0.0 | 19-26 |
| Ciprofloxacin (5 µg) | Fluoroquinolone | 0.0 ± 0.0 | 29-37 |
| Doxycycline (30 µg) | Tetracyclines | 0.0 ± 0.0 | - |
| Imipenem (10 µg) | Carbapenem | 7.3 ± 0.6 | 26-32 |
| Minimum inhibition concentration [MIC (µg/mL)] |  |  |  |
| Gentamicin | Aminoglycoside | 100 | 0.25-1 |
| Ciprofloxacin | Fluoroquinolone | 100 | 0.004-0.016 |
| Imipenem | Carbapenem | 0.19 | 0.06-0.5 |
| Meropenem | Carbapenem | 0.024 | 0.008-0.06 |
| Cephalexin | Beta-lactam | >100 | 4-16 |
| Tetracycline | Tetracycline | 100 | - |
| Amoxicillin | Beta-lactam | >100 | 2-8 |
| Amoxicillin-clavulanate | Beta-lactamase inhibitor | 50 | 2-8 |
| Colistin | Polymyxin | 0.19 | 0.25-2 |
| Erythromycin | Macrolide | >100 | - |

### 16S rRNA-Based Species Identification and Molecular Strain Typing

To confirm the identity of the bacterial isolate, the 16S rRNA gene was extracted using the “Extractseq” tool on the Galaxy platform and queried against the NCBI rRNA/ITS database using BLAST. This analysis confirmed the genus- and species-level identity of the isolate as *Escherichia coli*. The 16S rRNA gene sequence was submitted to GenBank under accession number PQ809548. MLST identified the isolate as sequence type 131 (ST131) (see Supplementary Material). *E. coli* ST131 is a globally disseminated, multidrug-resistant pathogen responsible for a high proportion of extraintestinal infections, particularly urinary tract infections (UTI) and bloodstream infections. It belongs to phylogenetic group B2 and is typically associated with the O25b:H4 serotype [45,46].

### Comprehensive Analysis of AMR Genes and Functional Annotations in the Genome of *E. coli* strain S3

The complete genome of *E. coli* strain S3 spans 5,332,658 bp and comprises five contigs linked to NCBI accession numbers CP196491 to CP196495 (Table 2). The primary chromosome is represented by Contig 1 (5,111,124 bp), while Contigs 2 to 5 correspond to plasmid-associated sequences. The circularity of both chromosome and plasmid elements was confirmed.

**Table 2:**
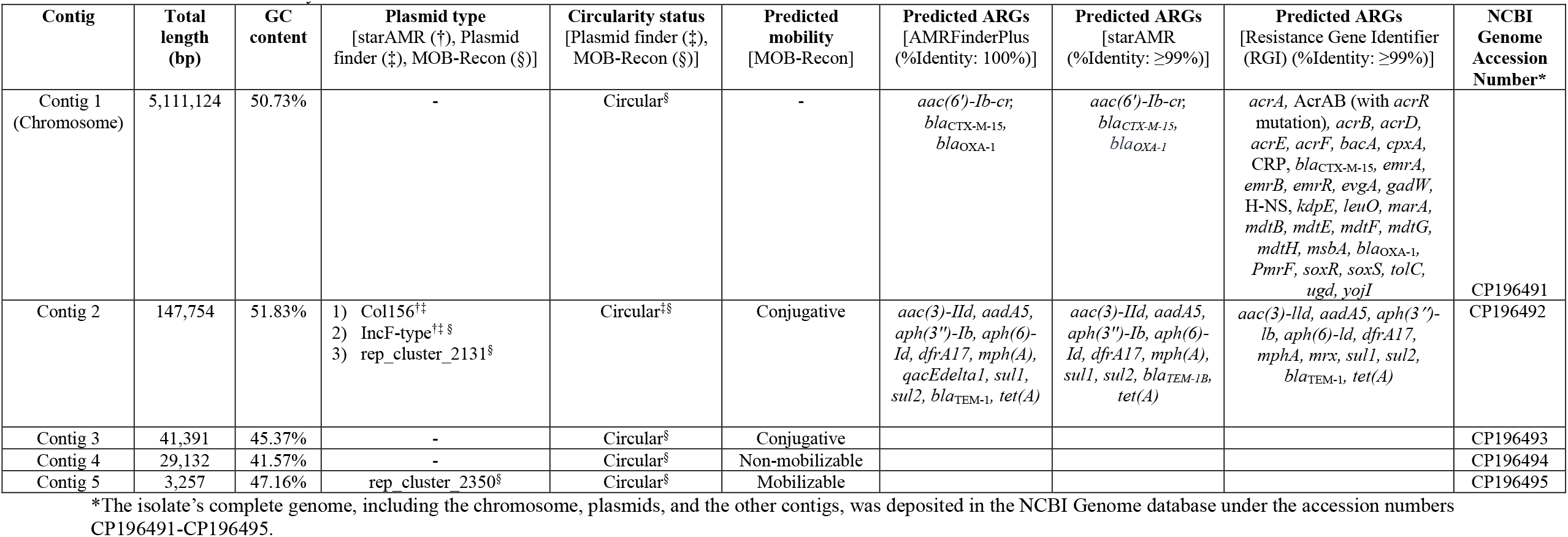
Genome assembly metrics and ARG content in *E. coli* strain S3.

| Contig | Total length (bp) | GC content | Plasmid type<br>[starAMR (†), Plasmid finder (‡), MOB-Recon (§)] | Circularity status<br>[Plasmid finder (‡), MOB-Recon (§)] | Predicted mobility<br>[MOB-Recon] | Predicted ARGs<br>[AMRFinderPlus (%Identity: 100%)] | Predicted ARGs<br>[starAMR (%Identity: ≥99%)] | Predicted ARGs<br>[Resistance Gene Identifier (RGI) (%Identity: ≥99%)] | NCBI Genome Accession Number* |
| --- | --- | --- | --- | --- | --- | --- | --- | --- | --- |
| Contig 1<br>(Chromosome) | 5,111,124 | 50.73% | - | Circular<sup>§</sup> | - | *aac(6')-Ib-cr*,<br>*bla<sub>CTX-M-15</sub>*,<br>*bla<sub>OXA-1</sub>* | *aac(6')-Ib-cr*,<br>*bla<sub>CTX-M-15</sub>*,<br>*bla<sub>OXA-1</sub>* | *acrA*, AcrAB (with *acrR* mutation), *acrB*, *acrD*, *acrE*, *acrF*, *bacA*, *cpxA*, CRP, *bla<sub>CTX-M-15</sub>*, *emrA*, *emrB*, *emrR*, *evgA*, *gadW*, H-NS, *kdpE*, *leuO*, *marA*, *mdtB*, *mdtE*, *mdtF*, *mdtG*, *mdtH*, *msbA*, *bla<sub>OXA-1</sub>*, *PmrF*, *soxR*, *soxS*, *tolC*, *ugd*, *yojI* | CP196491 |
| Contig 2 | 147,754 | 51.83% | 1) Col156<sup>†‡</sup><br>2) IncF-type<sup>†‡§</sup><br>3) rep_cluster_2131<sup>§</sup> | Circular<sup>‡§</sup> | Conjugative | *aac(3)-IId*, *aadA5*,<br>*aph(3'')-Ib*, *aph(6)-Id*, *dfrA17*, *mph(A)*,<br>*qacEdelta1*, *sul1*,<br>*sul2*, *bla<sub>TEM-1</sub>*, *tet(A)* | *aac(3)-IId*, *aadA5*,<br>*aph(3'')-Ib*, *aph(6)-Id*, *dfrA17*, *mph(A)*,<br>*sul1*, *sul2*, *bla<sub>TEM-1B</sub>*,<br>*tet(A)* | *aac(3)-IId*, *aadA5*, *aph(3'')-lb*, *aph(6)-Id*, *dfrA17*, *mphA*, *mrx*, *sul1*, *sul2*, *bla<sub>TEM-1</sub>*, *tet(A)* | CP196492 |
| Contig 3 | 41,391 | 45.37% | - | Circular<sup>§</sup> | Conjugative |  |  |  | CP196493 |
| Contig 4 | 29,132 | 41.57% | - | Circular<sup>§</sup> | Non-mobilizable |  |  |  | CP196494 |
| Contig 5 | 3,257 | 47.16% | rep_cluster_2350<sup>§</sup> | Circular<sup>§</sup> | Mobilizable |  |  |  | CP196495 |

\*The isolate's complete genome, including the chromosome, plasmids, and the other contigs, was deposited in the NCBI Genome database under the accession numbers CP196491-CP196495.

Contig 2 (147,754 bp) harbors multiple conjugative or mobilizable plasmid replicons and uncharacterized plasmid elements, including Col156 identified by PlasmidFinder and starAMR, rep_cluster_2131 detected by MOB-Recon, and IncF-type elements identified by all three plasmid identification tools, indicating a complex plasmid architecture. Contig 5 (3,257 bp) contains rep_cluster_2350, whereas Contigs 3 (41,391 bp) and 4 (29,132 bp) lack identifiable plasmid elements.

The labels rep_cluster_2131 and rep_cluster_2350 do not refer to traditional, specific plasmid replicon types such as Col156 or IncF-type plasmids. Instead, they are identifiers generated by the MOB-suite software package, representing clusters of plasmids grouped by overall sequence similarity and corresponding to broader evolutionary or taxonomic units rather than specific replication families or conserved replication initiation protein sequences [43].

Comprehensive resistome analysis using starAMR, AMRFinderPlus, and RGI at ≥99% identity threshold revealed an MDR profile for *E. coli* strain S3. In Contig 1, AMRFinderPlus and starAMR each detected three chromosomally borne, clinically significant ARGs associated with multiple antibiotic classes: *aac(6’)-Ib-cr* (aminoglycosides and fluoroquinolones), *bla*_CTX-M-15_(cephalosporins), *bla*_*OXA-1*_ (penicillins). RGI also identified *bla*_CTX-M-15_ and *bla*_OXA-1_ but additionally detected thirty ARGs not reported by the other tools. This broader detection included efflux and transport-associated genes such as *acrA, acrB, acrD, acrE, acrF, emrA, emrB, mdtB, mdtE, mdtF, mdtG, mdtH, tolC* and *yojI*, including the AcrAB-TolC efflux system with an *acrR* mutation. RGI also identified transcriptional regulators including *emrR, marA, soxS, soxR, cpxA, evgA*, CRP, *gadW*, H-NS, and *leuO*, as well as β-lactamases (*bla*_CTX-M-15_ and *bla*_OXA-1_) and cell envelope or membrane modification genes (*bacA, pmrF* and *ugd*).

Analysis of Contig 2 using all three tools revealed a wide range of ARGs spanning several antibiotic classes. While *qacEdelta1* (quaternary ammonium compounds) was detected only by AMRFinderPlus and *mrx* (macrolides) only by RGI, all tools consistently detected several ARGs, including *aac(3)-IId, aadA5, aph(3’’)-Ib* and *aph(6)-Id* (aminoglycosides), *dfrA17* (trimethoprim), *mph(A)* and *mrx* (macrolides), *sul1 and sul2* (sulfonamides), *bla*_TEM-1_ (penicillins), and *tet(A)* (tetracyclines).

The broader detection by RGI reflects its use of the Comprehensive Antibiotic Resistance Database, which includes a wide range of resistance determinants such as efflux pumps, regulatory genes, and intrinsic mechanisms. Its model-based approach captures genes that may contribute to resistance under specific conditions. In contrast, AMRFinderPlus and starAMR rely on more conservative, curated databases that prioritize well characterized, clinically relevant acquired resistance genes. Consequently, RGI provides a more expansive but less specific resistome profile, whereas the other tools yield more stringent, clinically focused results [40,41].

No ARGs were detected in Contigs 3, 4, or 5 by any of the tools, despite their circular structure and the presence of rep_cluster_2350 in Contig 5. The absence of ARGs at 99 to 100% identity thresholds suggests that these contigs likely serve functions unrelated to antimicrobial resistance. Overall, these findings demonstrate the presence of diverse and co localized ARGs in *E. coli* strain S3, highlighting its role as a reservoir for horizontally transferable resistance determinants mediated by complex plasmid structures.

### Virulence gene landscape indicates a high-risk, multi-pathotype *E. coli* strain

Table 3 summarizes the virulence gene profile of *E. coli* strain S3, revealing a dense repertoire of primarily chromosomally encoded determinants associated with adhesion, iron acquisition, and toxin production in Contig 1. Among the four plasmid-associated contigs (Contigs 2 - 5), only one virulence factor was identified, *senB*, which encodes an enterotoxin that induces intestinal fluid secretion and contributes to diarrheal disease.

**Table 3:** Virulence Gene Profile of *E. coli* strain S3.

| Contig | Gene | % Coverage | %Identity (≥99%) | Product | Implication for the isolate's virulence/pathogenicity |
| --- | --- | --- | --- | --- | --- |
| Contig 1 (chromosome) | *papB* | 100 | 99.36 | Regulatory protein PapB | Regulates expression of P fimbriae, which are critical for adhesion to uroepithelial cells, enhancing urinary tract infection (UTI) pathogenicity |
|  | *papC* | 100 | 99.88 | Usher protein PapC | Facilitates assembly and surface presentation of P fimbriae, promoting bacterial adhesion to host tissues, particularly in UTIs |
|  | *papD* | 100 | 100 | Chaperone protein PapD | Assists in folding and transport of P fimbriae subunits, essential for fimbrial biogenesis and host adhesion |
|  | *papJ* | 100 | 100 | P pilus assembly protein PapJ | Contributes to P fimbriae assembly, aiding in bacterial colonization of host tissues |
|  | *papK* | 100 | 100 | P pilus minor subunit PapK | Modulates fimbrial length and adhesion specificity, enhancing tissue attachment in UTIs |
|  | *papF* | 100 | 99.25 | P pilus minor subunit PapF | Critical for initiating fimbrial assembly and adhesion to host cells, increasing UTI virulence |
|  | *papG* | 100 | 99.11 | P pilus tip adhesin PapG | Mediates specific binding to Gal(α1-4)Gal receptors in the urinary tract, crucial for UTI pathogenesis |
|  | *papX* | 100 | 100 | PapX protein regulates flagellum synthesis to repress motility | Represses bacterial motility to enhance biofilm formation and tissue colonization, increasing pathogenicity |
|  | *yagW/ecpD* | 100 | 99.21 | Polymerized tip adhesin of ECP fibers | Promotes adhesion to intestinal epithelial cells, aiding in intestinal colonization and potential enteropathogenicity |
|  | *ykgK/ecpR* | 100 | 99.49 | Regulator protein EcpR | Regulates expression of *E. coli* common pilus (ECP), enhancing adhesion and biofilm formation in intestinal infections |
|  | *fes* | 100 | 99 | Enterobactin/ferric enterobactin esterase | Hydrolyzes enterobactin to release iron, supporting bacterial growth under iron-limited conditions in the host |
|  | *fepC* | 100 | 99.14 | Ferrienterobactin ABC transporter ATPase | Facilitates iron uptake via enterobactin, critical for bacterial survival and virulence in iron-scarce environments. Indirectly supports resistance by promoting fitness. |
|  | *entB* | 100 | 99.42 | Isochorismatase | Involved in enterobactin synthesis, enhancing iron acquisition and virulence |
|  | *ybtS* | 100 | 99.85 | Salicylate synthase Irp9 | Initiates yersiniabactin synthesis, a siderophore critical for iron acquisition in host tissues, enhancing virulence |
|  | *ybtX* | 100 | 99.69 | Putative signal transducer | Potentially regulates yersiniabactin system, supporting iron uptake and virulence |
|  | *ybtQ* | 100 | 99.56 | Inner membrane ABC-transporter YbtQ | Transports yersiniabactin, aiding iron acquisition and virulence |
|  | *ybtP* | 100 | 99.61 | Lipoprotein inner membrane ABC-transporter | Facilitates yersiniabactin transport, critical for iron uptake and pathogenesis |
|  | *ybtA* | 100 | 99.27 | Transcriptional regulator YbtA | Regulates yersiniabactin operon, enhancing iron acquisition and virulence |
|  | *irp2* | 100 | 99.44 | Yersiniabactin biosynthetic protein Irp2 | Essential for yersiniabactin synthesis, supporting iron uptake and virulence |
|  | *irp1* | 100 | 99.76 | Yersiniabactin biosynthetic protein Irp1 | Key in yersiniabactin production, critical for iron acquisition and pathogenesis |
|  | *ybtU* | 100 | 99.73 | Yersiniabactin biosynthetic protein YbtU | Supports yersiniabactin synthesis, aiding iron uptake and virulence |
|  | *ybtT* | 100 | 99.63 | Yersiniabactin biosynthetic protein YbtT | Contributes to yersiniabactin synthesis, enhancing virulence via iron acquisition |
|  | *ybtE* | 100 | 99.81 | Yersiniabactin siderophore biosynthetic protein | Initiates yersiniabactin biosynthesis, critical for iron uptake and virulence |
|  | *fyuA* | 100 | 99.8 | Pesticin/yersiniabactin receptor protein | Receptor for yersiniabactin, facilitating iron uptake and enhancing virulence. Indirect resistance role through survival |
|  | *iucA* | 100 | 99.83 | Aerobactin synthesis protein IucA | Involved in aerobactin synthesis, a siderophore enhancing iron uptake and virulence, particularly in extraintestinal infections |
|  | *iucB* | 100 | 99.79 | Aerobactin synthesis protein IucB | Supports aerobactin production, aiding iron acquisition and virulence. |
|  | *sat* | 100 | 99.56 | Secreted autotransporter toxin | Damages host cells, particularly in the urinary tract, increasing tissue invasion and UTI severity |
|  | *chuS* | 100 | 99.03 | Heme oxygenase ChuS | Degrades heme to release iron, supporting bacterial growth in blood-rich environments |
|  | *chuA* | 100 | 99.54 | Outer membrane heme/hemoglobin receptor ChuA | Binds heme/hemoglobin, critical for iron acquisition in blood, enhancing systemic infection virulence |
|  | *chuT* | 100 | 99.8 | Periplasmic heme-binding protein ChuT | Transports heme across the periplasm, supporting iron uptake and virulence |
|  | *chuX* | 100 | 100 | Putative heme-binding protein ChuX | Likely aids heme utilization, enhancing virulence in blood-rich environments |
|  | *chuU* | 100 | 99.5 | Heme permease protein ChuU | Facilitates heme transport across the inner membrane, supporting virulence |
|  | *chuV* | 100 | 99.75 | ATP-binding hydrophilic protein ChuV | Provides energy for heme transport, critical for virulence in systemic infections |
| Contig 2 | *senB* | 99.91 | 99.74 | Enterotoxin | Causes fluid secretion in the intestine, contributing to diarrheal disease |
| Contig 3 | - | - | - | - |  |
| Contig 4 | - | - | - | - |  |
| Contig 5 | - | - | - | - |  |

The strain’s virulence landscape is dominated by a complete *pap* operon, consistent with a uropathogenic phenotype driven by efficient urinary tract adhesion and colonization. Multiple siderophore systems, including enterobactin, yersiniabactin, and aerobactin, along with heme uptake genes, indicate strong iron-scavenging capacity that enhances survival and invasiveness in iron-limited host environments. Notably, the presence of the plasmid-borne *senB* enterotoxin broadens the clinical potential of the strain to include enteric disease, highlighting the role of mobile genetic elements in the dissemination of clinically relevant virulence traits in poultry settings.

## DISCUSSION

Building on our recent investigation of AMR in poultry farms in Enugu, Nigeria, the present study characterizes an MDR *E. coli* strain identified during that surveillance [32]. While our earlier work provided a broad overview of AMR trends in the region, this follow-up analysis focuses on *E. coli* strain S3, which exhibits a notably alarming resistance profile, offering new insights into the mechanisms and potential public health implications of emerging MDR bacteria in agricultural settings. By examining this uniquely resistant isolate, we provide a detailed analysis of its genetic determinants of multidrug resistance and virulence factors, complementing the broader trends reported in our earlier work [32, 33,34].

### Phenotypic AMR profiling of *E. coli* strain S3

The phenotypic AMR profile of the *E. coli* strain S3 reveals a concerning level of multidrug resistance, with implications for both public health and agricultural antibiotic stewardship. Notably, *E. coli* strain S3 displayed resistance with no inhibition zone formation to multiple first-line and critically important antibiotics, including cefotaxime, ampicillin, erythromycin, gentamicin, ciprofloxacin, and doxycycline. This extensive resistance was further corroborated by MIC testing, which showed extremely high MIC values (≥100 µg/mL) for gentamicin, ciprofloxacin, tetracycline, cephalexin, erythromycin, and amoxicillin - well above clinical breakpoints for susceptibility [35]. Such a broad-spectrum resistance phenotype is consistent with previous reports of multidrug-resistant (MDR) *E. coli* in poultry environments, where extensive antibiotic use has promoted resistance. For instance, Abreu *et al*. [3] reviewed the status and innovative strategies for controlling AMR in poultry production, Aworh *et al*. [25] documented the genetic relatedness of MDR *E. coli* from humans, chickens, and poultry environments in Nigeria, and Hedman *et al*. [47] reported high levels of AMR in poultry farms in low-resource settings.

In Nigeria, antimicrobial use in poultry is widespread, often prophylactic, with drugs obtained over-the-counter [32]. The most used classes include tetracyclines, penicillins, aminoglycosides, fluoroquinolones, macrolides, and sulphonamides [32]. Surveys indicate routine administration of these antibiotics for prevention, treatment, and growth promotion [32,25]. Correspondingly, poultry-associated isolates frequently show high resistance to these same classes; for instance, tetracycline, sulphonamide, and fluoroquinolone resistance rates often exceed 50% [34,48]. The resistance profile of our isolate aligns with these trends, reflecting selective pressure from locally prevalent antimicrobial practices and highlighting the relevance of national AMR surveillance gaps in interpreting local resistance dynamics [32,34].

### Genomic Determinants of AMR in *E. coli* strain S3

*E. coli* strain S3 exhibits a resistance profile that aligns with broader patterns of AMR reported in poultry production systems. Racewicz et al. [48] documented a high prevalence of AMR genes, including integron-associated elements, in *E. coli* isolates from poultry production systems, highlighting the genetic mechanisms underpinning resistance dissemination. Similarly, Veloo et al. [49] reported widespread MDR *E. coli* in both broiler and indigenous farm settings in Malaysia, underscoring the potential for such environments to act as reservoirs of resistant strains. Together, these findings suggest that the resistance profile observed in strain S3 in this current study reflects a broader trend of escalating AMR within intensive and semi-intensive poultry production systems [21,47], reinforcing the need for strengthened surveillance and antimicrobial stewardship interventions. The complete genome of *E. coli* strain S3, assembled into five contigs spanning 5.33 Mb, reflects the genomic complexity characteristic of MDR *E. coli* strains.

The complete genome of strain S3, assembled into five contigs spanning 5.33 Mb, demonstrates the genomic complexity typical of MDR *E. coli*. The presence of a primary chromosome (Contig 1; 5.11 Mb; GC content 50.73%) alongside multiple conjugative or mobilizable plasmid-associated contigs highlights the strain’s capacity for horizontal gene transfer (HGT) and genomic plasticity [4,5,6,17,50]. Notably, the strain harbors a diverse set of plasmid replicons, including Col156, IncF type plasmids, and rep_cluster_2131 and rep_cluster_2350 within a single genome. This combination reflects substantial genetic flexibility and an enhanced ability to acquire and disseminate AMR and other adaptive traits [51].

Col156 is a small mobilizable plasmid commonly associated with *qnrS* genes and fluoroquinolone resistance in both avian and clinical isolates [52]. IncF-type plasmids are well established contributors to the global spread of multidrug resistance in both human and animal settings [6,48,51]. Although less frequently reported, rep_cluster_2131 has been identified in *E. coli* isolated from humans, wild animals, and companion animals. Similarly, rep_cluster_2350 has been reported in *E. coli* isolated from humans, livestock, wild animals, food, companion animals, and environmental sources [53]. The coexistence of these conjugative or mobilizable elements within a single strain illustrates the complexity of resistance gene exchange in agroecosystems and highlights the zoonotic potential of poultry-derived *E. coli* [16,23].

Chromosomally encoded ARGs, including *aac(6’)-Ib-cr, bla*_CTX-M-15_, and *bla*_OXA-1_, further strengthen the resistance profile of strain S3. These genes, often linked to IncF plasmids and globally disseminated MDR lineages, are commonly mobilized by insertion sequences such as IS*Ecp1* or transposons, enabling their transfer from plasmids to the chromosome. Once integrated, they are stably inherited during cell division, even in the absence of antibiotic pressure, unlike plasmids which may be lost if they impose a fitness cost [51,55]. The presence of *bla*_CTX-M-15_ and *bla*_OXA-1_ is particularly concerning due to their association with ESBL production and oxacillinase-mediated resistance, both of which are linked to treatment failure in serious infections [54]

In addition to chromosomal determinants, numerous plasmid-borne ARGs contribute to the MDR phenotype. These include genes conferring resistance to aminoglycosides [*aac(3)-IId, aadA5, aph(3’’)-Ib*, and *aph(6)-Id,Ib*], trimethoprim [*dfrA17*], macrolides [*mph(A)* and *mrx*], sulfonamides [sul1/sul2], β-lactams [*bla*_TEM-1_] and tetracyclines [*tet(A)*]. These genes are disseminated through both vertical transmission and HGT, particularly via conjugation, while transposable elements facilitate their movement between plasmids and the chromosome. The co localization of multiple ARGs on single mobile elements promotes co selection, allowing exposure to one antimicrobial agent to maintain resistance to several others [18,56].

No ARGs were detected at a ≥99% identity threshold in the circular uncharacterized plasmids located in contigs 3 and 4 or in rep_cluster_2350 within contig 5. These elements may instead contribute to plasmid stability, host fitness, or virulence. This supports the view that some plasmids function as genetic scaffolds for future ARG acquisition or aid bacterial adaptation through non-resistance mechanisms [56,57].

The MDR phenotype of strain S3, particularly its resistance to beta lactams, fluoroquinolones, and aminoglycosides, highlights the risk of ARG dissemination through HGT in poultry production systems in Nigeria and reflects global concerns regarding antibiotic use in food animal production [9,58]. This study provides the first whole genome-based characterization of plasmid-mediated multidrug resistance in poultry-associated *E. coli* strain S3 from Enugu, Nigeria. It offers important insights into the local resistome and mobilome and establishes a foundation for ongoing genomic surveillance within the poultry production chain.

### One Health Implications of MDR *E. coli* ST131 in Poultry: Virulence, Persistence, and Environmental Dissemination

Analysis of *E. coli* strain S3 identified multiple virulence-associated genes involved in adhesion, iron acquisition, and toxin production (Table 3), suggesting an enhanced capacity for host colonization and persistence. However, the presence of these genes alone does not confirm clinical pathogenicity. The pap gene cluster (*papB–D, F, G, J, K, X*) encodes P fimbriae, structures that facilitate adhesion to uroepithelial cells and contribute to biofilm formation [59]. Specific adhesin variants such as *papGII* and *papGIII* have been linked to distinct UTI syndromes in clinical isolates, but their detection in strain S3 indicates only pathogenic potential rather than confirmed involvement in UTI [59-61]. Regulatory genes such as *papB* and *papX* may further enhance persistence by modulating fimbrial expression and motility under host stress conditions [60,62].

The strain also harbored multiple iron acquisition systems, including enterobactin (*fes, fepC, entB*), yersiniabactin (*ybt* genes, *fyuA*), aerobactin (*iucAB*), and heme uptake (*chu* genes), all of which enhance bacterial survival in iron-limited environments such as the bloodstream, urinary tract, intestinal tract, respiratory mucosa, and within macrophages and neutrophils [63-66]. These siderophore systems are frequently associated with extraintestinal pathogenic *E. coli* (ExPEC) and avian pathogenic *E. coli* (APEC), supporting adaptation to extraintestinal niches, although their presence does not establish disease causation [67,68]. They may nevertheless promote persistence under antimicrobial stress, indirectly supporting AMR. In addition, toxin-associated genes such as sat, encoding an autotransporter toxin, and *senB*, encoding the ShET2 enterotoxin, may contribute to tissue damage in urinary and intestinal environments [69,70]. Adhesion-related genes *yagW/ecpD* and *ykgK/ecpR*, which encode *E. coli* common pili, may further enhance colonization and biofilm formation, thereby improving bacterial fitness and persistence without directly conferring AMR [70,71].

Taken together, the virulence-associated gene repertoire of strain S3 suggests an ExPEC-like profile characterized by adhesins, siderophores, and toxins that may enhance extraintestinal survival, biofilm-mediated tolerance, and persistence under antimicrobial pressure. Based on established ExPEC criteria, the isolate can therefore be tentatively classified as an ExPEC-like strain [72,73]. However, this classification is based solely on genomic content and does not confirm pathogenicity in the absence of clinical infection data.

The isolation of MDR *E. coli* ST131 from chicken droppings has important implications for public health, veterinary medicine, and environmental management within the One Health framework. ST131 is a globally disseminated multidrug-resistant lineage, commonly associated with phylogenetic group B2 and serotype O25b:H4, and is a major cause of extraintestinal infections such as UTIs and bloodstream infections in humans [45,46]. Its detection in poultry waste suggests that poultry may serve as a reservoir for clinically relevant antimicrobial resistant lineages, challenging the view of ST131 as a predominantly human-associated pathogen [74,75].

The identified ST131 isolate, confirmed through 16S rRNA sequencing and MLST, possessed an ExPEC-like virulence profile that included P fimbriae, multiple siderophore systems, and toxin-associated genes. These factors enhance adhesion, iron acquisition, biofilm formation, and persistence under stress, all of which may support long-term survival and the maintenance of AMR [67,76]. Although these virulence determinants do not confirm pathogenicity, they indicate a strong potential for extraintestinal survival and infection, particularly under antimicrobial selection pressure.

From a public health perspective, the detection of ST131 in poultry environments raises significant zoonotic concerns. Poultry-associated ST131 sublineages may enter the human food chain through contaminated meat or environmental exposure, colonize the human gastrointestinal tract, and later cause infections such as UTIs or urosepsis [74,75]. The public health threat is compounded by the frequent presence of resistance determinants such as ESBL genes and colistin resistance genes, which can compromise treatment options and increase the risk of therapeutic failure [74, 77, 78]. In addition, poultry workers may become asymptomatic carriers through occupational exposure, facilitating silent dissemination within the community [78,79].

In veterinary medicine, ST131 may function as avian pathogenic *E. coli* (APEC), contributing to colibacillosis, respiratory disease, pericarditis, and systemic infections in poultry. These infections are associated with increased mortality, particularly in young chicks, and may be perpetuated through vertical transmission from breeder hens to offspring. Such persistence within flocks can lead to chronic infections, reduced feed efficiency, carcass condemnation, and substantial economic losses [80,81]. The virulence-associated genes identified in the isolate, including adhesins, siderophores, and toxins, likely enhance colonization and survival in avian hosts, increasing the difficulty of disease control [81-83].

The environmental implications are equally concerning. Poultry droppings, particularly when used untreated as organic fertilizer, provide a route for ST131 and associated AMR genes to enter soil, water, and crop systems [84-86]. The environmental resilience of this lineage allows it to persist outside the host, creating long-term reservoirs of AMR in the ecosystem. This reservoir may facilitate indirect transmission to wildlife and humans, expanding the ecological and epidemiological reach of the pathogen [87-90]. The fecal-environmental cycle therefore plays an important role in the maintenance and dissemination of ST131 beyond farm settings [91,92].

These findings strongly support the One Health concept, emphasizing the interconnectedness of human, animal, and environmental health [93-95]. The detection of MDR ST131 in poultry waste demonstrates the need for integrated surveillance systems that extend beyond clinical settings to include agricultural and environmental reservoirs. It also underscores the importance of antimicrobial stewardship and strengthened farm biosecurity to reduce the emergence, persistence, and spread of high-risk antimicrobial-resistant pathogens.

In summary, the isolation of *E. coli* ST131 from chicken droppings is not merely a microbiological observation but evidence of a multidimensional public health challenge involving zoonotic transmission, veterinary disease burden, and environmental dissemination of antimicrobial resistance. Its presence in poultry waste highlights the need for coordinated interventions across public health, veterinary, and environmental sectors to interrupt transmission pathways and mitigate the global spread of this high-risk lineage.

### Limitations of the Study

This study focuses on an important MDR *E. coli* isolate representing a unique MDR phenotype from a poultry farm. The isolate was selected for the genomic analyses reported in this paper because of its distinct resistance profile, as determined by disk diffusion and broth dilution antibiotic susceptibility testing. Focusing on a single strain may limit the generalizability of our findings to the broader population of poultry-associated *E. coli* in Enugu or across Nigeria. However, despite this limitation, the study provides critical insights into the genomic architecture and zoonotic potential of MDR *E. coli* from Nigerian poultry, thereby establishing a foundation for future surveillance and functional investigations.

## CONCLUSION

This study provides a whole-genome characterization of the MDR *E. coli* strain S3 isolated from Nigerian poultry, highlighting its resistance and virulence potential. Phenotypic AMR profiling showed complete resistance to multiple critically important antibiotics, including aminoglycosides, fluoroquinolones, β-lactams, macrolides, and tetracyclines, while remaining susceptible to carbapenems and colistin. Genomic analysis identified chromosomal ARGs, including *aac(6’)-Ib-cr*, which confers resistance to aminoglycosides and fluoroquinolones; *bla*_CTX-M-15_, associated with cephalosporin resistance; and *bla*_OXA-1_, linked to penicillin resistance. In addition, multiple efflux and transport-associated genes, transcriptional regulators, and cell envelope or membrane modification genes were detected, reflecting a broad AMR profile. Several plasmid-borne ARGs, including genes conferring resistance to aminoglycosides, trimethoprim, macrolides, sulfonamides, penicillins, tetracyclines, were also identified. The presence of multiple plasmids, including Col156 and IncF-type plasmids, indicates a strong potential for horizontal gene transfer. Furthermore, virulence genes associated with adhesion, iron acquisition, and toxin production suggest a notable pathogenic risk. MLST classified the strain as ST131. From a One Health perspective, the isolation of this strain from chicken droppings highlights a multidimensional public health challenge involving zoonotic transmission, veterinary disease burden, and environmental dissemination of AMR. The detection of this high-risk sequence type in poultry waste underscores the need for integrated surveillance systems that extend beyond clinical settings to include agricultural and environmental reservoirs. It also emphasizes the importance of antimicrobial stewardship and strengthened farm biosecurity to reduce the emergence, persistence, and spread of high-risk antimicrobial-resistant pathogens. Overall, this study demonstrates the potential for poultry farms to serve as reservoirs of MDR pathogens and reinforces the need for prudent antibiotic use, enhanced biosecurity, and comprehensive surveillance.

## DECLARATIONS

### Ethics approval and consent to participate

Ethical approval for the study was granted by the Enugu State Ministry of Health (Reference number: MH/MSD/REC21/630).

### Consent for publication

Not applicable

### Competing interests

The authors declare no competing interests.

### Funding

This work was supported by the National Institutes of Health (NIH) through the Combatting Antimicrobial Resistance in Africa Using Data Science (CAMRA) grant (project number 5U54TW012056-03).

### Authors’ Contributions

Conceptualization: CPE and PME. Sample collection and field investigation: CPE, CE, PME, EAN. Bioinformatics and data analysis: CPE, PME, CT, CJC. Interpretation of results: CPE, PME, EAN, MUA, CE. Writing – original draft: CPE, PME, MUA. Writing – review and editing: CPE, CE, PME, EAN, CT, CJC, MUA. All authors read and approved the final manuscript.

## Acknowledgements

The authors gratefully acknowledge Enugu State University of Science and Technology (ESUT), Enugu, Nigeria, for administrative support provided during this project.

## Data availability

Whole genome sequencing data generated from Illumina-hybrid assemblies have been deposited in the NCBI database under BioProject ID PRJNA1293944, BioSample ID SAMN50084471 and GenBank accession numbers CP196491-CP196495. The corresponding 16S rRNA gene sequence has been submitted to GenBank under accession number PQ809548.

## Supplementary material

The growth of *E. coli* strain 3 on Coliform ChromoSelect Agar (CCA), the MALDI-TOF identification results, the AMR analyses using AMRFinderPlus, starAMR, and RGI, the bacterial 16S rRNA sequence query in NCBI, and the virulence gene profile analysis using ABRicate are available in the supplementary materials.

## Notes

https://www.ncbi.nlm.nih.gov/nuccore/2873623433?log$=activity

### Competing Interest Statement

The authors have declared no competing interest.

### Summary of Updates

This current version of the manuscript is the most updated.

